# High-light adaptation via ferredoxin-mediated tuning of the photosynthesis–photoprotection trade-off

**DOI:** 10.64898/2026.06.18.733257

**Authors:** Julián Gabriel Bultri, Candela Brugnara, Celina Lobais, Michael Melzer, Nicolás E Blanco

**Affiliations:** Centre of Photosynthetic and Biochemical Studies (CEFOBI-CONICET), Faculty of Biochemical Science and Pharmacy, Rosario National University, S2002LRK, Rosario, Argentina; Leibniz-Institut für Pflanzengenetik und Kulturpflanzenforschung, Physiologie und Zellbiologie, 06466 Gatersleben, Germany

**Keywords:** *Nicotiana tabacum*, photosynthesis, photoprotection, high-light, Ferredoxin, PSI-acceptor side limitation

## Abstract

Plants continuously adjust photosynthesis to balance growth and photoprotection under changing environmental conditions. Environmental fluctuations frequently impose a mismatch between energy production and CO2 assimilation. How photochemical reactions are regulated to maintain performance under these conditions remains a central question in plant biology.
We previously developed transplastomic tobacco (*Fd1-*OE plants) overexpressing ferredoxin (Fd) displaying enhanced photoprotection and growth penalties with a variegated leaf phenotype under greenhouse conditions. Here, we investigate how these plants respond to different growth irradiances using physiological, ultrastructural, and photosynthetic analyses, including PAM, gas exchange, and P700 absorbance measurements, and dynamic-light assays.
*Fd1*-OE plants progressively recovered growth, leaf phenotype and photosynthetic performance as growth irradiance increased, reaching near WT performance at 1400 μmol m⁻² s⁻¹. This enhanced adaptation to “high-light” was associated with a larger fraction of open PSII reaction centers and enhanced NPQ. Dynamic-light analyses further revealed faster plastoquinone (PQ) turnover, a more oxidized PQ pool and enhanced electron withdrawal downstream of PSI.
Our results indicate that Fd overexpression redefines the balance between photochemistry and photoprotection. This adjustment shifts adaptation toward higher irradiance and enhances photosynthetic performance under changing light environments. Electron partitioning downstream of PSI emerges as a promising target to improve photosynthetic resilience.

**One sentence summary:** Overexpression of Fd1 in tobacco plants adjusts photosynthesis/photoprotection trade off to enhance high-light adaptation

## Introduction

Photosynthetic electron transport continuously adjusts to fluctuations in irradiance to maintain the balance between energy capture and metabolic consumption (Huner *et al*., 1998; Johnson, 2025). This balance depends on the coordination between light-driven ATP and NADPH production and their downstream consumption by primary metabolism, mainly CO2 assimilation. Environmental constraints that reduce assimilation rates or stromal metabolism frequently disrupt this balance (Huner *et al*., 2016; Becklin *et al*., 2021). For instance, factors that limit CO2 assimilation and/or Calvin-Benson-Bassham (CBB) cycle activity in illuminated leaves (e.g., cold temperature) reduce NADPH consumption, promoting stromal over-reduction. Under these conditions, continued light absorption and electron flow impose additional pressure on the photosynthetic electron transport chain (PETC), as redox turnover of its components becomes increasingly restricted. Failure to dissipate excess excitation energy and associated electron pressure increases the risk of photo-oxidative damage and photoinhibition (Aro *et al*., 1993; Gjindali & Johnson, 2023). To provide operational flexibility, a set of photoprotective mechanisms function alongside the PETC to dissipate or redistribute excess absorbed light energy and modulate electron fluxes (Walker et al., 2020; Gjindali & Johnson, 2023). Because these mechanisms often operate through energy dissipation or electron redistribution, they define a trade-off with photosynthesis efficiency. Therefore, maintaining photostasis in a context of fluctuating environmental conditions requires a fine tuning between photochemical reactions and photoprotective processes (Huner et al., 2012).

Photoprotective mechanisms operate within both spatial and temporal hierarchies coordinated by sensing redox imbalances within the PETC. Among these mechanisms, non-photochemical quenching (NPQ), photosynthetic control, and alternative electron flows are coordinately activated when electron transport exceeds stromal electron sink capacity (Shikanai & Yamamoto, 2017; Ruban & Wilson, 2021; Schansker, 2022). Hence, the buildup of proton motive force, particularly its ΔpH component, constitutes a central trigger coordinating these three mechanisms (Kramer *et al*., 2003; Tikhonov, 2015). At the same time, both Photosystem I (PSI) activity and the redox poise of the plastoquinone (PQ) pool are major regulatory nodes for adjusting PETC function to external conditions, although they are rarely considered within an integrated framework (Alric, 2015; Tikhonov & Vershubskii, 2017). In this context, PSI acceptor-side limitation, defined as restricted availability of oxidized stromal electron acceptors capable of sustaining electron withdrawal from PSI, represents a common consequence of adverse growth conditions. The resulting imbalances constraints photosynthetic efficiency and increases the risk of PSI photodamage (Suorsa *et al*., 2012; Chaux *et al*., 2015; Tikhonov, 2015).

Several genetic strategies have been explored to alleviate PSI acceptor-side limitation while simultaneously sustaining photosynthetic efficiency. One straightforward rationale has been to overexpress proteins associated with stromal electron partitioning and alternative electron flows. However, overexpression of proteins involved in PSI regulation, alternative electron flows or electron partitioning (e.g., PGR5, flavodiiron proteins, and ferredoxins), frequently results in complex physiological trade-offs rather than straightforward improvements in photosynthetic performance (Yamamoto *et al*., 2006; Okegawa *et al*., 2007; Blanco *et al*., 2013; Yamamoto *et al*., 2016; Wada *et al*., 2018; Rizzetto *et al*., 2025). Interestingly, in some cases, the establishment of a leaf-variegated phenotype indicates perturbations in photosynthetic regulation together with penalties in growth and chloroplast function (Okegawa *et al*., 2007; Rosso *et al*., 2009; Okegawa *et al*., 2010). Thus, variegation illustrates that balancing photochemistry and photoprotection requires tight regulation and ultimately determines the permissive limits for plant growth. Together, these observations highlight that changes in electron partitioning downstream of PSI can reshape photosynthetic regulation, yet their consequences for plant acclimation remain largely unexplored. Previously, we characterized *Fd1*-OE plants, which also display a variegated phenotype associated with enhanced NPQ and growth penalties under greenhouse conditions (Blanco et al., 2013). Here, we investigated whether this altered photosynthesis–photoprotection trade-off modifies the adaptive response and acclimation of plants to changing light environments. Unlike classical variegated mutants, higher growth irradiance progressively restored wild-type growth, photosynthetic performance, and leaf phenotype in *Fd1*-OE plants. This unusual behavior respect to other variegated mutants prompted us to evaluate how the elevated Fd1 levels modify the regulation of photosynthetic electron transport and thereby the adaptive and acclimation response.

## Materials and Methods

### Plant Growth Conditions

For all the experiments, wild-type (WT) *Nicotiana tabacum* (cv. Petit Havana) and ferredoxin overexpressing Fd1-OE lines under the same background (Blanco *et al*., 2013) were grown in a controlled environment chamber under a 16/8 light/dark photoperiod at 25°C and 60% relative humidity. Seedlings were kept at 350 µmol m⁻² s⁻¹ for a period of 2 weeks, until developing 2 fully-expanded leaves. Then, they were transferred to individual pots and, after 1 week, assigned a corresponding final light intensity (ranging from 30 to 1400 µmol m⁻² s⁻¹). 5-week-old plants were used for the experiments. The maximum irradiance value of 1400 µmol m⁻² s⁻¹ was selected based on light intensities typically recorded under outdoor conditions around 9:00 AM during mid-November in Rosario, Argentina (32.97029° S, 60.62414° W).

### Sucrose Supplementation

Seeds were surface-sterilized by incubation in a 1:10 (v/v) sodium hypochlorite solution for 10 min and rinsed four times with sterile distilled water. Sterile seeds were sown on Petri dishes containing Murashige and Skoog (MS) medium supplemented with 0%, 1%, or 3% (w/v) sucrose. Sucrose was sterilized by filtration through a 0.2 µm membrane filter and added to the medium prior to solidification. Plates were maintained under the same controlled conditions described above and grown at 150 µmol m⁻² s⁻¹ for 20 days.

### Measurements of Chlorophyll Fluorescence

Plants were dark-adapted for a minimum of 20 minutes prior to analysis, and measurements were taken from the first fully expanded leaf. Baseline fluorescence in the dark (F0) was obtained under a very low-intensity measuring beam (620 nm; 0.1 µmol m⁻² s⁻¹). A saturating pulse (500 ms, 8000 µmol m⁻² s⁻¹) was then applied to determine the maximum fluorescence in darkness (Fm) and again during exposure to actinic light (AL) to obtain Fm′. For each AL step, the steady-state fluorescence level (Fs) was recorded. From these values, the maximum PSII efficiency (*Fv*/*Fm*), with *F*v=*Fm*-*F0,* and non-photochemical quenching, NPQ = (*Fm-Fm′)*/*Fm′*, were calculated. The effective quantum yield of PSII, Y(II), was determined as *(Fm′*-*Fs*)/*Fm*′. The parameter *qL*, representing the fraction of open PSII reaction centers and the redox state of the plastoquinone pool, was calculated as (*Fm*′-*Fs*)/(*Fm*′-*F0*′) × (*F0*′/*Fs*), where *F0*′ was derived from *F0*/(*Fv*/*Fm*+*F0*/*Fm*′). P700 redox dynamics were monitored via absorbance changes at 830 and 875 nm. The maximal oxidation level of P700 in darkness (Pm) was induced by applying a saturating pulse in the presence of far-red background light (720 nm). The corresponding maximal level during AL exposure (Pm′) was determined with an additional saturating pulse, and the steady-state P700⁺ signal (P) was recorded immediately beforehand. PSI quantum yields were calculated as follows: Y(I) = (Pm′−P)/Pm; acceptor-side limitation, Y(NA) = (Pm-Pm′)/Pm; and donor-side limitation, Y(ND) = P/Pm. These three yields satisfy Y(I)+Y(NA)+Y(ND) = 1.

Chlorophyll fluorescence measurements were performed using a PAR-FluorPen FP 110/D (PSI), and a LI-6800 photosynthesis system (LI-COR Biosciences). Combined chlorophyll fluorescence and P700 absorption analyses were conducted with a DUAL-PAM-100 system (Heinz Walz), using ten biological replicates for each condition.

To record the OJIP transient, the saturating pulse was set to 3000 µmol m⁻² s⁻¹ with a duration of 800 ms, ensuring full closure of PSII reaction centers. Fluorescence emission was collected at high-frequency sampling throughout the induction period, capturing the characteristic O–J–I–P rise. *F0*, *Fm* and intermediate phases (Fj and Fi) were extracted automatically by the LI-6800 software using default detection thresholds. For each genotype and light treatment, 6 biological replicates were analyzed.

For NPQ induction, dark-adapted fluorescence parameters were obtained, and then the leaves were illuminated with actinic light at 2000 µmol m⁻² s⁻¹ to induce the maximal heat-dissipative response. NPQ induction was monitored for 10 minutes, with saturating pulses delivered every 20 seconds. Following illumination, leaves were returned to darkness to record NPQ relaxation. Fluorescence was monitored for 15 minutes, using saturating pulses applied at progressively longer intervals to capture the full decay of the quenched state. All measurements were performed on the first fully expanded leaf of at least 6 biological replicates.

The rapid light response of the photosynthetic apparatus was assessed by exposing intact leaves to a stepwise series of actinic light intensities (0, 100, 150, 250, 350, 450, 750, 1150, 1450, 2150, 3050, and 3650 µmol m⁻² s⁻¹). For each light step, leaves were allowed to reach a steady fluorescence signal before applying a saturating pulse. The saturating pulses were used to determine the fluorescence parameters following the formulas described above.

After obtaining dark-adapted and steady-state parameters, plants growing at 1400 µmol m⁻² s⁻¹ were exposed to a fluctuating light stress treatment corresponding to 4 minutes of low light (62 µmol m⁻² s⁻¹) followed by 1 minute of high-light intensity (1961 µmol m⁻² s⁻¹). After four repetitions, the profile of every parameter was analyzed.

### Gas Exchange Measurements

CO2 assimilation and chlorophyll fluorescence were recorded simultaneously using a LI-6800 gas-exchange system paired with the 6800-01A fluorometer head (LI-COR Biosciences). Prior to data collection, leaves were allowed to reach steady-state under the following reference conditions: 420 ppm CO2, 1500 µmol m⁻² s⁻¹ irradiance, 27 °C leaf temperature, 65% relative humidity, and a flow rate of 500 µmol s-1. Light-response curves were generated by keeping the reference CO2 concentration fixed at 420 ppm and gradually decreasing the actinic light from 2000 µmol m⁻² s⁻¹ to darkness in 3-minute decrements. Actinic illumination consisted of a red/blue mixture (90%/10%) for all measurements. 10 biological replicates were analyzed for each condition.

### Pigment Content Analysis

Leaf pigment levels were quantified using ethanol extracts from standardized leaf tissue. Three discs (8 mm diameter) were collected from the tip region of the first fully expanded leaf, weighed, and incubated in 1 mL of 96% ethanol for 48 h in complete darkness. Absorbance of the extracts was recorded at 470, 648.1, and 664.1 nm using a Synergy HTX MultiMode Reader (BioTek), adapting the equations from (Lichtenthaler & Buschmann, 2001) for chlorophyll and carotenoid determination. Readings at 750 nm were used to address non-specific turbidity or background absorbance unrelated to pigment content. Six biological replicates were analyzed.

### Western Blotting

Soluble proteins were extracted from 200 mg of leaf tissue collected from 5-week-old plants. Samples were immediately frozen in liquid nitrogen for homogenization in 1 ml of extraction buffer (50 mM Tris–HCl, pH 7.5; 1 mM EDTA; 5 mM MgCl2; 150 mM NaCl, and 1 mM PMSF). The total protein extract was obtained by cold centrifugation at 10,000 g for 1 min and stored in Sample buffer (2% SDS, 10% glycerol; 2.5% β-mercaptoethanol). 10 µg of protein were resolved by 12% SDS–polyacrylamide gel electrophoresis and transferred to a nitrocellulose membrane. Membranes were blocked with 2% (w/v) milk in TBS. For each rabbit antibody incubation, final concentrations were reached in 2% milk TBS solution, incubating overnight. Anti-rabbit IgG conjugated with horseradish peroxidase (1:20000, Cell Signaling) was used for chemiluminescence detection, using the SuperSignal West Femto Maximum Sensitivity Substrate (Thermo Scientific), and visualized after 1 minute of reaction using an Amersham Imager 600 (GE Healthcare Life Sciences). The following antibodies from Agrisera were used: anti-PsbA (AS05 084, 1:10000), anti-PetC (AS08 330, 1:10000), anti-PsaA (AS05 172, 1:5000), anti-Lhca1 (AS01 005, 1:5000), anti- Lhcb2 (AS01 003, 1:5000), anti-Lhcb2-P (AS13 2705, 1:10000), anti-Lhcb4 (AS04 045, 1:7000), anti-PGR5 (AS16 3985, 1:5000), anti-PGRL1 (AS19 4311, 1:1000), anti-NdhH (AS16 4065, 1:5000), anti-VDE (AS15 3091, 1:2000), anti-FDX1 (AS06 121, 1:1000).

### Histological and Ultrastructural Analysis of Leaf Tissue

Leaf samples of at least 3 different plants were collected from the first fully expanded leaves and cut into ∼1-2 mm2 sections and used for combined conventional and microwave-assisted fixation, dehydration and resin embedding as shown in Supplementary Table S1. Ultrathin sectioning and ultrastructure analysis were carried out as described previously (Schumann *et al*., 2017). Images were captured from mesophyll chloroplasts for each plant and light condition. Chloroplasts from 50 cells per genotype were imaged and used for analysis of grana number, thylakoid organization, and plastoglobuli characteristics.

### Statistical analysis

All data were analyzed using InfoStat software (v2020e). For experiments involving a single growth irradiance, or for comparison among genotypes within the same irradiance, differences were evaluated by one-way analysis of variance (ANOVA), with genotype as the main factor. When significant effects were detected, means were compared using Tukey’s honestly significant difference (HSD) test. For experiments involving multiple growth irradiances, data were analyzed by two-way ANOVA with genotype and growth irradiance as fixed factors, including their interaction term. When significant effects were detected, mean comparisons were performed using Tukey’s HSD test. Statistical significance was established at p ≤ 0.05.

## Results

### “Low-light” conditions and sugar supplementation do not restore the wild-type phenotype in Fd1-OE plants

Previous characterization of *Fd1*-OE plants revealed growth penalties, delayed development, and leaf variegation when plants were grown under “greenhouse conditions” (PPFD of 150 μmol m⁻² s⁻¹). Similar phenotypes in other mutants have been associated with imbalances in the redox poise of the PETC, particularly in the plastoquinone (PQ) pool, and were alleviated by reducing growth light intensities (Rosso *et al*., 2009; Foudree *et al*., 2010; Okegawa *et al*., 2010). To assess whether similar mechanisms underpin the *Fd1*-OE plant phenotype under “greenhouse conditions”, *Fd1*-OE and control (hereinafter named WT) plants were grown at lower irradiance (30 μmol m⁻² s⁻¹, also referred to as “low-light” conditions) (Fig. 1). Under reduced PPFD, *Fd1*-OE plants failed to recover the WT phenotype or growth rate, maintaining the variegation in leaves (Fig. 1a, insets). Following the same trend observed in control plants, reducing growth irradiance did not increase the photosynthetic pigment content in *Fd1*-OE plants but instead led to a decrease in chlorophyll and carotenoids (Table 1). These reductions were even more pronounced than those observed in WT siblings grown under the same “low-light” conditions. A second transplastomic line showed the same response to reduced growth light intensity, confirming the absence of transgene insertion effects in these plastome-transformed plants (Fig. S1) (Blanco *et al*., 2013; Bock, 2015).

**Figure 1.**
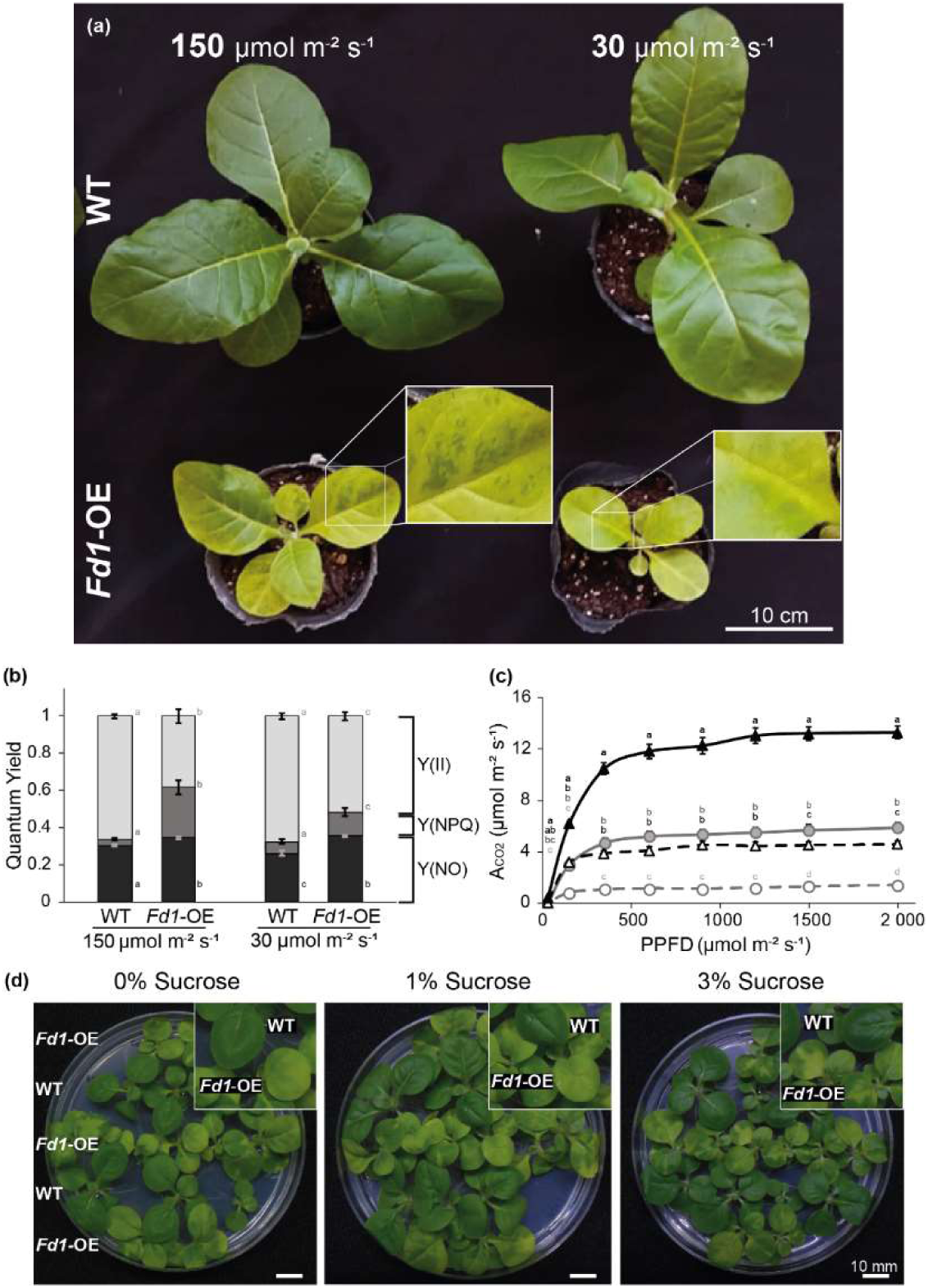
Low-light conditions and sugar supplementation do not restore the wild-type phenotype in *Fd1*-OE plants. *Fd1*-OE plant phenotype was evaluated under different growth conditions. (a) Five-week-old control (WT, top) and ferredoxin-overexpressing (*Fd1*-OE, bottom) plants grown under “greenhouse” (150 μmol m⁻² s⁻¹) and “low-light” (30 μmol m⁻² s⁻¹) conditions. (b) Steady-state PS II quantum yield parameters Y(II) (light grey), Y(NPQ) (dark grey), and Y(NO) (black), measured at the corresponding growth irradiance. (c) Light-response curves of CO₂ assimilation rates for WT (black triangles) and *Fd1*-OE plants (grey circles). Empty symbols and dashed lines correspond to plants grown under “low-light” conditions, whereas filled symbols and solid lines correspond to plants grown under “greenhouse” conditions. Data points represent means ± SE (n = 10). Different letters indicate significant differences between genotypes at a given light intensity (two-way ANOVA followed by Tukey’s multiple comparison test, p ≤ 0.05). (d) WT and *Fd1*-OE plants grown on MS agar plates supplemented with different sucrose concentrations under “greenhouse” conditions. D.

**Table 1.**
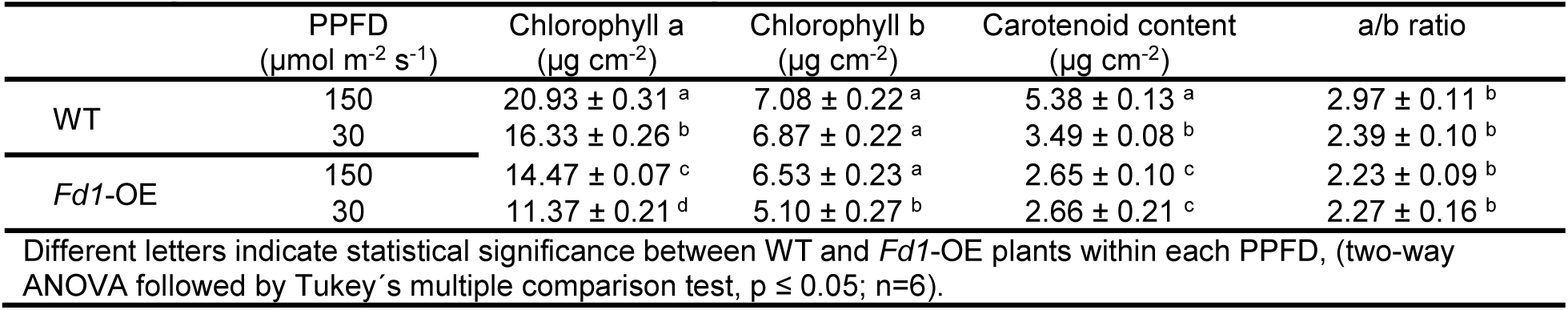
Pigment content in plants exposed to low light intensities.

| Table 1. Pigment content in plants exposed to low light intensities. |  |  |  |  |  |
| --- | --- | --- | --- | --- | --- |
| | PPFD<br>($\mu\text{mol m}^{-2} \text{s}^{-1}$) | Chlorophyll a<br>($\mu\text{g cm}^{-2}$) | Chlorophyll b<br>($\mu\text{g cm}^{-2}$) | Carotenoid content<br>($\mu\text{g cm}^{-2}$) | a/b ratio |
| WT | 150 | $20.93 \pm 0.31^a$ | $7.08 \pm 0.22^a$ | $5.38 \pm 0.13^a$ | $2.97 \pm 0.11^b$ |
| | 30 | $16.33 \pm 0.26^b$ | $6.87 \pm 0.22^a$ | $3.49 \pm 0.08^b$ | $2.39 \pm 0.10^b$ |
| *Fd1*-OE | 150 | $14.47 \pm 0.07^c$ | $6.53 \pm 0.23^a$ | $2.65 \pm 0.10^c$ | $2.23 \pm 0.09^b$ |
| | 30 | $11.37 \pm 0.21^d$ | $5.10 \pm 0.27^b$ | $2.66 \pm 0.21^c$ | $2.27 \pm 0.16^b$ |

Different letters indicate statistical significance between WT and *Fd1*-OE plants within each PPFD, (two-way ANOVA followed by Tukey's multiple comparison test, $p \leq 0.05$ ; $n=6$ ).

Consistent with the expected response to “low light” conditions, CO₂ assimilation rates were reduced in control and *Fd1*-OE plants, with the latter showing the lower values (Fig. 1c). The partitioning of absorbed energy followed the same trend for both genotypes (Fig. 1b). However, the photochemical usage of absorbed light of *Fd1*-OE plants remained lower than control plants due to a higher portion of this energy being dissipated in a non-photochemical way (Y(NO) plus Y(NPQ)).

Similarly, neither variegation nor reduced growth was restored to WT levels when *Fd1*-OE plants were grown with exogenous carbon supply (MS medium supplemented with 0%, 1%, or 3% sucrose) (Fig. 1d). Moreover, both lines exhibited delayed growth at 3% sucrose. Taken together, these results indicate that neither reducing irradiance nor exogenous carbon supply is sufficient to rescue the defective growth phenotype of *Fd1*-OE plants. These observations suggest that the overexpressing phenotype does not arise solely from excessive light stress or carbon limitation, but instead reflects broader Fd1-dependent alterations in photosynthetic regulation.

### Increases in growth light intensities restore growth and photosynthetic performance in Fd1-OE plants

The impaired growth rate, photosynthetic performance, and leaf-variegated phenotype of *Fd1*-OE plants under greenhouse conditions were exacerbated by reducing growth light intensity (PPFD from 150 to 30 μmol m⁻² s⁻¹), contrasting with other variegated mutants (Blanco *et al*., 2013). To gain insight into this alternative Fd-mediated photosynthetic regulation, *Fd1*-OE plants were grown under dynamic illumination regimes with peak PPFDs of 350, 600, 900 and 1400 μmol m⁻² s⁻¹, spanning a broad range irradiances simulating outdoor-light patterns (Fig. 2; Fig. S2). Increasing PPFD progressively shifted the *Fd1*-OE plant characteristics toward WT-like phenotype (Fig. 2a). At 1400 μmol m⁻² s⁻¹, *Fd1*-OE plants exhibited near WT-like appearance, pigmentation, and growth (Table 2). The Fd transgene expression was not affected by the increase in growth light intensity (Fig. 2a, inset). Maximum CO2 assimilation rates and maximum photosynthetic quantum yield (*Fv*/*Fm*) increased in parallel with growth PPFD in *Fd1*-OE plants, closely approaching wild-type values at the highest light intensity (Fig. 2b and c). The changes in *Fv*/*Fm* were associated with variations in both fluorescence parameters, *Fm* and *F*0.

**Figure 2.**
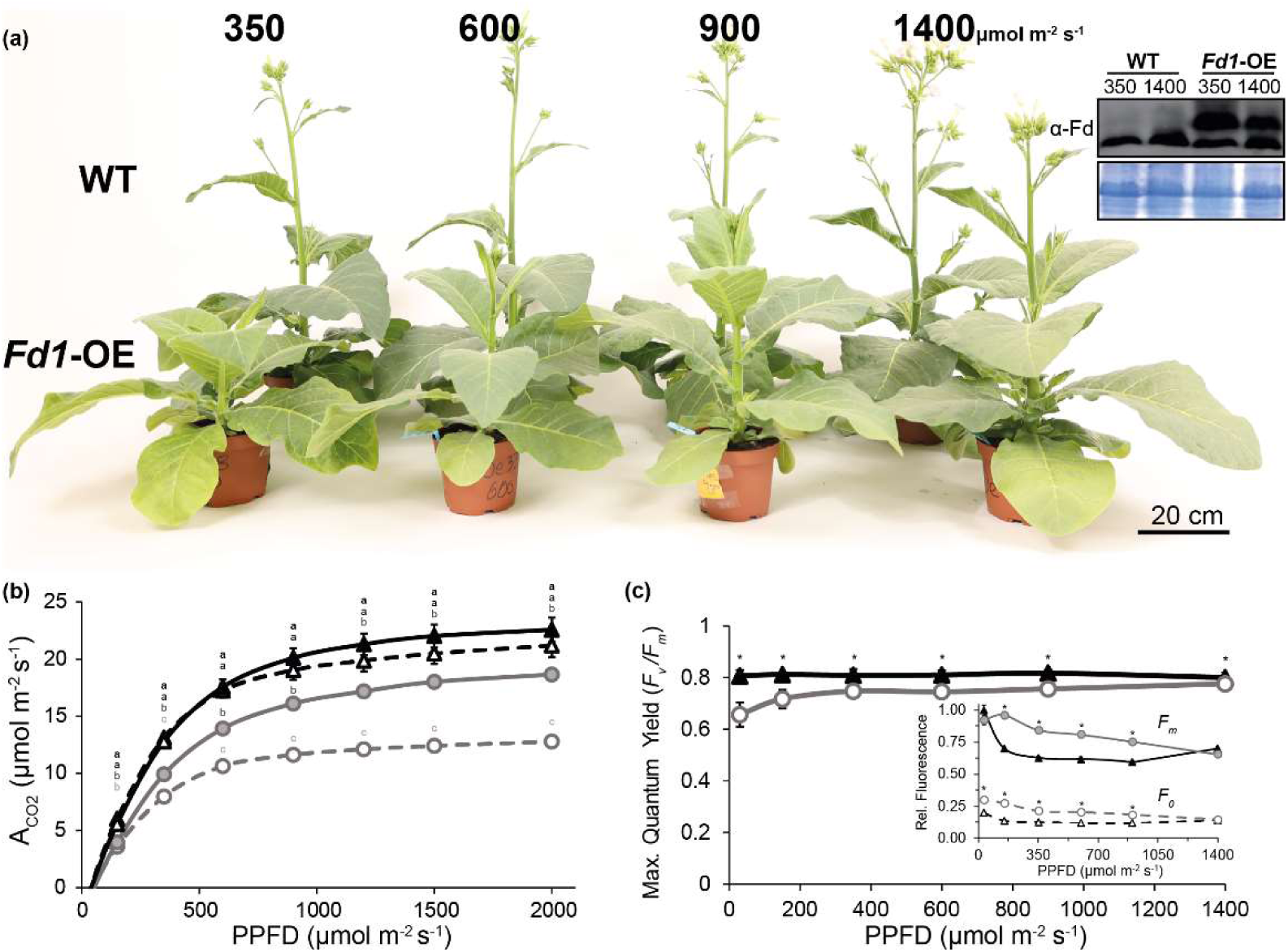
*Fd1*-OE plants recover an almost wild-type phenotype under outdoor-like growth conditions. (a) Seven-week-old WT (top) and *Fd1*-OE (bottom) plant phenotypes grown under increasing light intensities (350, 600, 900, 1400 μmol m-2 s-1, left to right), (b) Light-response curves of CO₂ assimilation rates for WT (black triangles) and *Fd1*-OE (grey circles) plants grown at 350 μmol m-2 s-1 (dashed lines, empty symbols) and 1400 μmol m-2 s-1 (solid lines and symbols), showing enhanced photosynthetic activity in *Fd1*-OE plants at the highest light intensity (c) Maximum PSII quantum yield (*Fv/Fm*) of WT and *Fd1*-OE plants for each PPFD evaluated, including parameters obtained from plants growing at PPFD of 30 and 150 μmol m^-2^ s^-1^ (Fig. 1). *Fd1*-OE plants presented maximum values of *Fv/Fm* parameter when grown at 1400 μmol m^-2^ s^-1^. The inset shows the corresponding F0 (dashed lines) and *Fm* (solid lines) values for each condition. Data points represent means ± SE (n = 20). Different letters in (b) indicate significant differences according to two-way ANOVA followed by Tukey’s multiple comparison test (p ≤ 0.05). Asterisks in (c) indicate significant differences between genotypes within each PPFD according to one-way ANOVA followed by Tukey’s multiple comparison test (p ≤ 0.05).

**Table 2.** Pigment content in plants exposed to increasing light intensities.

| | PPFD<br>($\mu\text{mol m}^{-2} \text{s}^{-1}$) | Chlorophyll a<br>($\mu\text{g cm}^{-2}$) | Chlorophyll b<br>($\mu\text{g cm}^{-2}$) | Carotenoid content<br>($\mu\text{g cm}^{-2}$) | a/b ratio |
| --- | --- | --- | --- | --- | --- |
| WT | 350 | $28.45 \pm 0.03^a$ | $7.19 \pm 0.22^a$ | $7.38 \pm 0.21^a$ | $3.96 \pm 0.05^a$ |
| | 1400 | $28.66 \pm 0.11^a$ | $6.34 \pm 0.22^b$ | $8.15 \pm 0.37^a$ | $4.53 \pm 0.11^b$ |
| *Fd1*-OE | 350 | $21.68 \pm 0.19^b$ | $7.12 \pm 0.23^b$ | $5.62 \pm 0.15^b$ | $3.06 \pm 0.11^c$ |
| | 1400 | $27.04 \pm 0.07^c$ | $7.26 \pm 0.27^b$ | $7.42 \pm 0.30^a$ | $3.75 \pm 0.14^a$ |

Different letters indicate statistical significance between WT and *Fd1*-OE plants within each PPFD, (two-way ANOVA followed by Tukey's multiple comparison test, $p \leq 0.05$ ; $n=6$ ).

Together, these results indicate that the phenotype and growth penalties associated with elevated Fd content in *Fd1*-OE plants are strongly dependent on growth light intensity and become progressively attenuated as growth irradiance increases. Among the tested conditions, the highest growth light intensity of 1400 μmol m⁻² s⁻¹ provided the most favorable growth conditions for *Fd1*-OE plants.

### Ultrastructural analysis of chloroplasts in Fd1-OE plants is highly sensitive to growth conditions

The striking changes in phenotype, maximum quantum yield and CO2 assimilation rates observed in *Fd1*-OE plants grown at 1400 μmol m⁻² s⁻¹ compared with 350 μmol m⁻² s⁻¹ prompted us to further examine photosynthetic performance through chloroplast structural organization. First, chloroplast ultrastructure of *Fd1*-OE and WT plants grown under “greenhouse” and “high-light” conditions was analyzed by transmission electron microscopy (TEM) (Fig. 3). In WT plants, chloroplasts displayed well-organized grana stacks under both growth conditions, with a moderate increase in plastoglobuli (PG) size and thicker grana with smaller diameters at higher light intensities. Such structural features have previously been observed in response to moderate changes in growth conditions (Schumann *et al*., 2017). In contrast, *Fd1*-OE plants again exhibited more pronounced differences between the two growth light conditions. Under “greenhouse conditions”, thylakoid membranes of *Fd1*-OE plants showed reduced grana height and an increased number of PG compared with WT counterparts. These differences were further accentuated under “high-light” conditions. When comparing growth conditions within the *Fd1*-OE genotype, thylakoid remodeling was characterized by more prominent stroma lamellae, markedly reduced grana appression, and larger and more abundant PG. Altogether, these results suggest that the phenotypic reversion of *Fd1*-OE plants grown at 1400 μmol m⁻² s⁻¹, including the loss of variegation, does not reflect the recovery of functionally WT chloroplasts but rather the establishment of an alternative structural organization. The similarities between the thylakoid membrane remodeling in *Fd1*-OE chloroplasts and the differences in chloroplast ultrastructure between fully developed shade and sun leaves prompted us to evaluate the components of the photosynthetic electron transport chain (Shuang *et al*., 2022).

**Figure 3.**
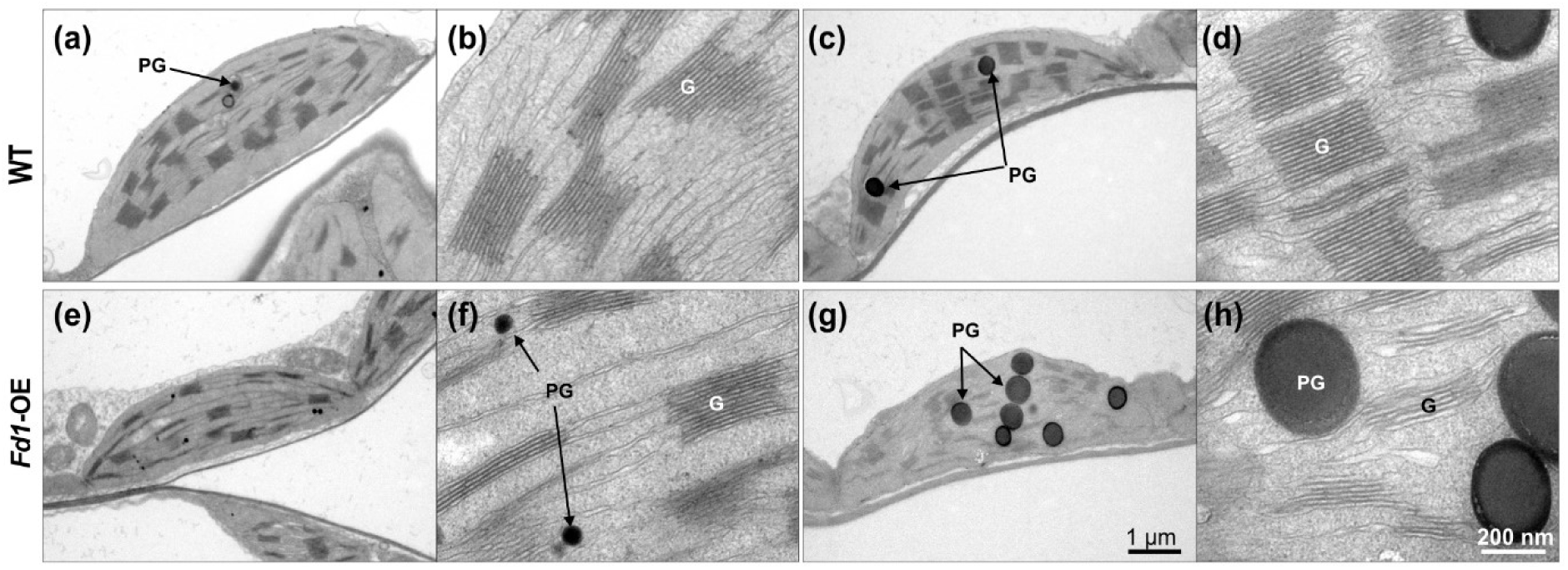
Ultrastructure of chloroplasts in *Fd1*-OE plants shows profound remodeling of thylakoid membranes at “high-light” conditions. The phenotype of *Fd1*-OE plants was tested against different growth conditions. Transmission electron microscopy (TEM) images of chloroplasts from each genotype (WT, top and *Fd1*-OE, bottom) grown under “greenhouse“ (a, b, e, f) and “high-light” (c, d, g, h) conditions. Stacked grana are visible as darker grey structures, plastoglobuli (PG) can be seen as black dots. Differences in thylakoid organization and plastoglobuli accumulation illustrate the structural response to light intensity and genetic background.

### Fd1-OE plants exhibit a Fd-dependent alternative regulation of photosynthesis

In light of the reorganization of the thylakoid membranes and the associated adjustment of photosynthesis to growth conditions (Lichtenthaler *et al*., 1981; Johnson & Wientjes, 2020), we analyzed the abundance of major PETC components and the steady-state photosynthetic performance (Fig. 4 and Table 3). Consistent with the stable expression of the heterologous Fd1 in *Fd1*-OE plants, the abundance of core PETC components did not differ between genotypes or across growth conditions (Fig. 4a). Proteins analyzed included PSII (PsbA), cytochrome *b6f* (PetC), PSI (PsaA), and light-harvesting complex (LHC) proteins (Lhca1, Lhcb2 and Lhcb4). Similarly, the abundance of regulatory proteins involved in alternative electron flow pathways (PGR5, PGRL1, NdhH) or non-photochemical quenching (VDE) remained unchanged. The only detectable difference between WT and *Fd1*-OE plants was a lower phosphorylation level of Light-harvesting chlorophyll a/b-binding protein 2 (Lhcb2) in transgenic plants. Because Lhcb2 phosphorylation participates in regulating state transitions and influences the partitioning of LHCII between photosystems, the lower phosphorylation status observed in *Fd1*-OE plants may reflect differences in excitation energy distribution, and be aligned with the observed thylakoid membrane remodeling (Longoni *et al*., 2015; Longoni & Goldschmidt-Clermont, 2021). However, neither LHCII distribution nor photosynthetic supercomplex organization was directly assessed in this study. Overall, altered photosynthetic behavior in *Fd1*-OE plants was not associated with major compositional remodeling of the PETC.

**Figure 4.**
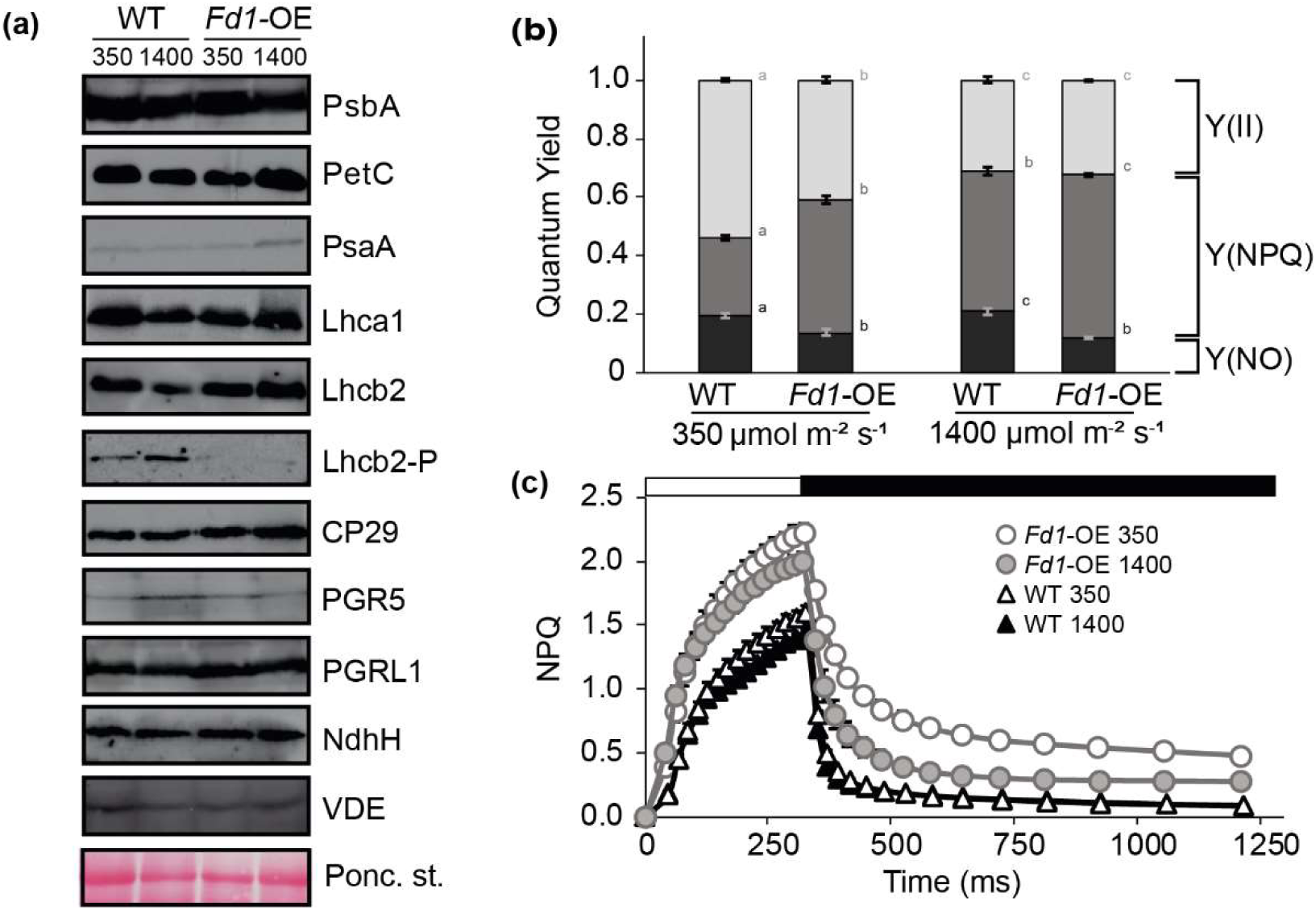
Fd1 overexpression results in a differential regulation of photosynthetic performance under steady-state conditions. (a) Analysis of the composition of the PETC by immunoblot analysis of essential subunits in *Fd*1-OE and WT plants grown at 350 and 1400 μmol m⁻² s⁻¹. Loading control obtained by Ponceau staining are shown below the membranes. (b) Absorbed energy partitioning of *Fd1*-OE and WT plants at different growth light intensity reveals a more stable balance between photochemical (Y(II)) and dissipative reactions. (c) NPQ induction curves show that the higher NPQ in *Fd1*-OE plants is sustained across different growth light intensities. Data represent means ± SE (n ≥ 10). Different letters indicate significant differences between WT and *Fd1-*OE plants within each PPFD (two-way ANOVA followed by Tukeýs multiple comparison test, p ≤ 0.05).

**Table 3.**
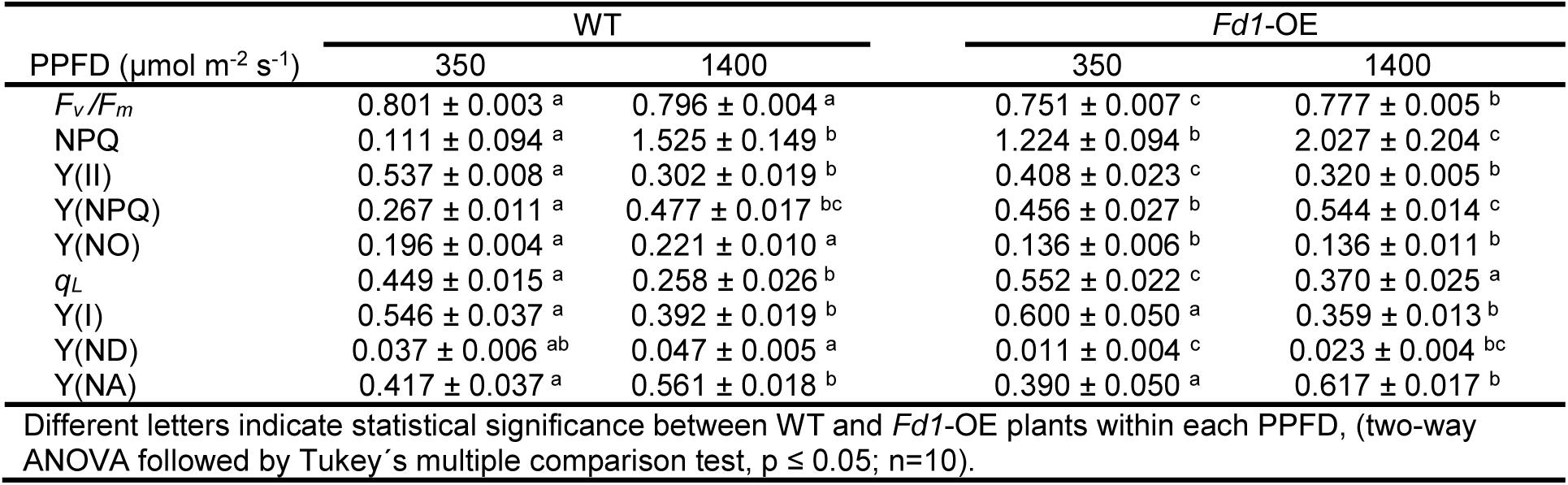
Steady-state chlorophyll fluorescence parameters of WT and Fd1-OE plants growing under “high-light” conditions.

Under “low-light” and “greenhouse” conditions, *Fd1*-OE plants exhibited enhanced NPQ, Y(NPQ), and Y(NO) at the expense of photochemical efficiency (Y(II)) (Fig. 1b) (Blanco et al., 2013). When grown at 350 and 1400 μmol m⁻² s⁻¹, *Fd1*-OE plants maintained higher NPQ than WT (Fig. 4b and c, Table 3). However, the increase in Y(NPQ) from “low” to “high-light” was markedly attenuated in *Fd1*-OE compared with WT plants (∼19% Y(NPQ)*Fd1*-OE1400-*Fd1*-OE350, vs ∼78% Y(NPQ)WT1400-WT350). Notably, WT plants grown at 1400 μmol m⁻² s⁻¹ reached Y(NPQ) values similar to those in *Fd1*-OE plants grown at lower irradiance (i.e., 350 μmol m⁻² s⁻¹).

This more moderate increase in energy dissipation (Y(NPQ)) in *Fd1*-OE plants at 1400 μmol m⁻² s⁻¹ was accompanied by a smaller reduction in the photochemical component Y(II) (Fig. 4b and Table 3). Consequently, the balance between photochemical energy use (Y(II)) and photoprotective dissipation (Y(NPQ)) across growth irradiances was more stable in *Fd1*-OE plants. As a result, PSII photochemical capacity (*qL*) was higher in *Fd1-*OE than in WT plants at 1400 μmol m⁻² s⁻¹ (*qL*_WT1400 ∼0.26 vs *qL*_*Fd1*-OE1400 ∼0.37; Table 3).

In contrast, PSI-related steady-state parameters showed comparatively minor differences between genotypes (Table 3). Parameters associated with PSI activity (Y(I), Y(NA), Y(ND)) were largely unaffected, although a reduction in PSI donor-side limitation (Y(ND)) was observed in *Fd1*-OE plants grown at “high-light” conditions (Y(ND)*Fd1*-OE 1400 ∼ 0.023 vs Y(ND)*Fd1*-OE 350 = 0.011), a response not observed in WT plants.

Taken together, these results indicate that *Fd1*-OE plants exhibit an alternative photosynthetic regulation that is not driven by changes in PETC composition but rather by modifications in energy partitioning and electron transport regulation. Under “high-light” conditions, *Fd1*-OE plants sustain photoprotective processes while maintaining higher PSII photochemical capacity, suggesting an altered regulation of redox poise within the electron transport chain.

### Fd1 tunes photosynthetic induction and acclimation to dynamic light environments

The differences in steady-state photosynthetic parameters across different growth light intensities, and the disappearance of the conditional “variegated” phenotype in *Fd1*-OE plants grown at 1400 μmol m⁻² s⁻¹ suggest an alternative photosynthetic regulation associated with elevated Fd1 content. Moreover, the observed *Fd1*-OE phenotype shares similarities with “high-light” adaptive responses (Walker *et al*., 2020; Gjindali & Johnson, 2023). To evaluate whether this differential adaptive response also influences photosynthetic regulation under changing irradiance, regulation of photosynthesis was studied through OJIP curves, analysis of the post-illumination chlorophyll fluorescence transient rise (PIFR), and rapid light curves (RLC) (Fig. 5 and 6).

**Figure 5.**
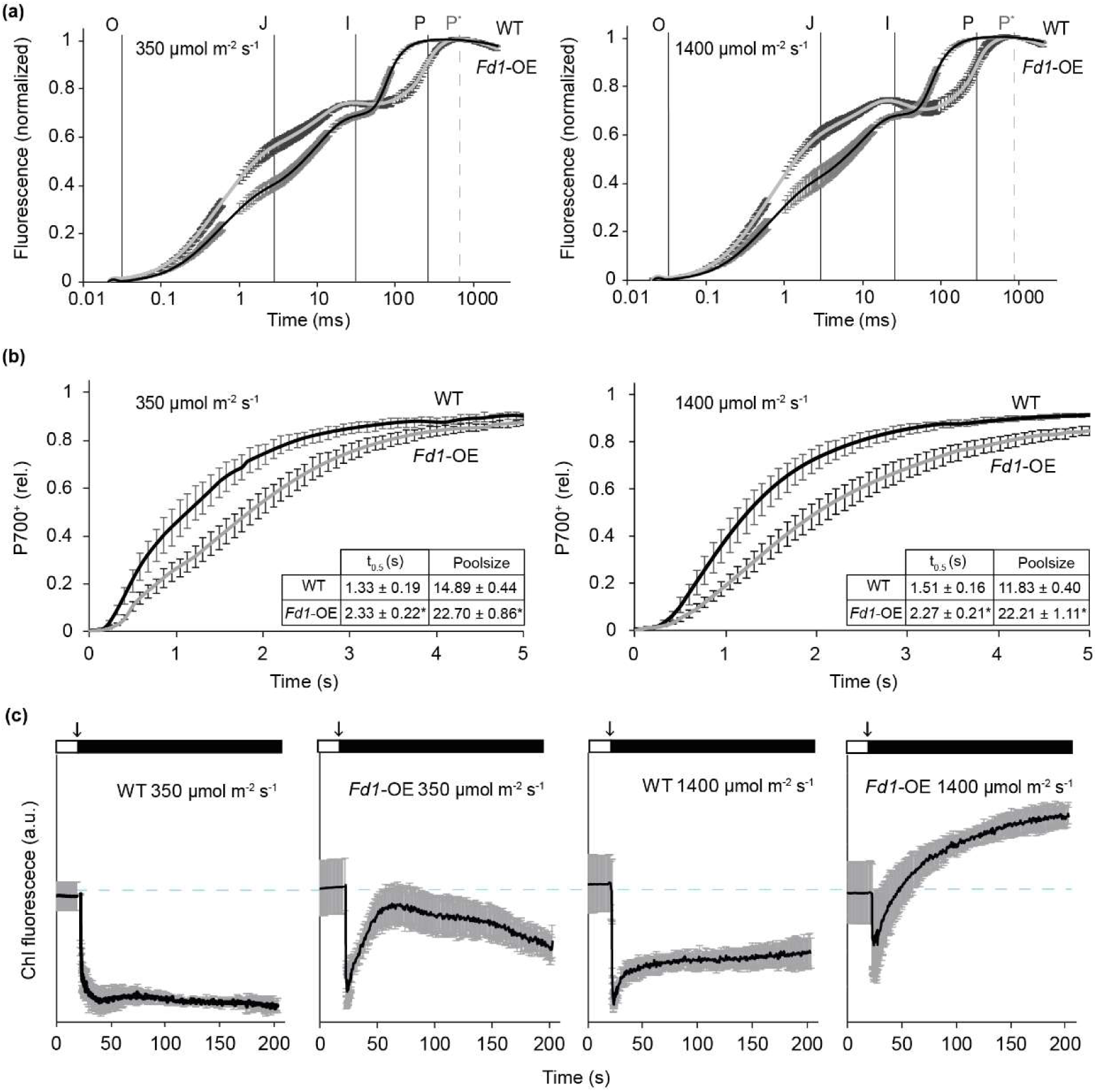
*Fd1*-OE plants exhibited a differential regulation of the photosynthesis during light transitions. (a) Polyphasic chlorophyll a fluorescence induction (OJIP) curves for *Fd1*-OE and WT plants grown at 350 and 1400 µmol m-2 s-1, revealing differential redox kinetics alongside PETD for *Fd1*-OE plants. Data were double-normalized at O and P levels. The O-J, J-I and I-P phases are defined, and plotted on a logarithmic time scale from 10 μs to 1 s. The delayed P peak observed in *Fd1*-OE plants is indicated by dotted grey lines (P*). (b) Post-illumination chlorophyll fluorescence transient rise (PIFR) curves measured for each growth condition after exposure to the corresponding growth irradiance for at least 12 min, followed by transfer to darkness. (c) PSI re-oxidation kinetics measured under far-red illumination following a multiple-turnover flash. The inset shows re-oxidation parameters derived from at least 6 biological replicates (means ± SE). Half-time (t₀.₅) corresponds to the time required to reach 50% of the re-oxidation amplitude, whereas the area above the curve (Poolsize parameter) is related to the amount of reduced electron carriers, predominantly within the plastoquinone pool. Asterisks in (c) indicate significant differences between genotypes within each growth irradiance (one-way ANOVA followed by Tukey’s multiple comparison test, p ≤ 0.05).

**Figure 6.**
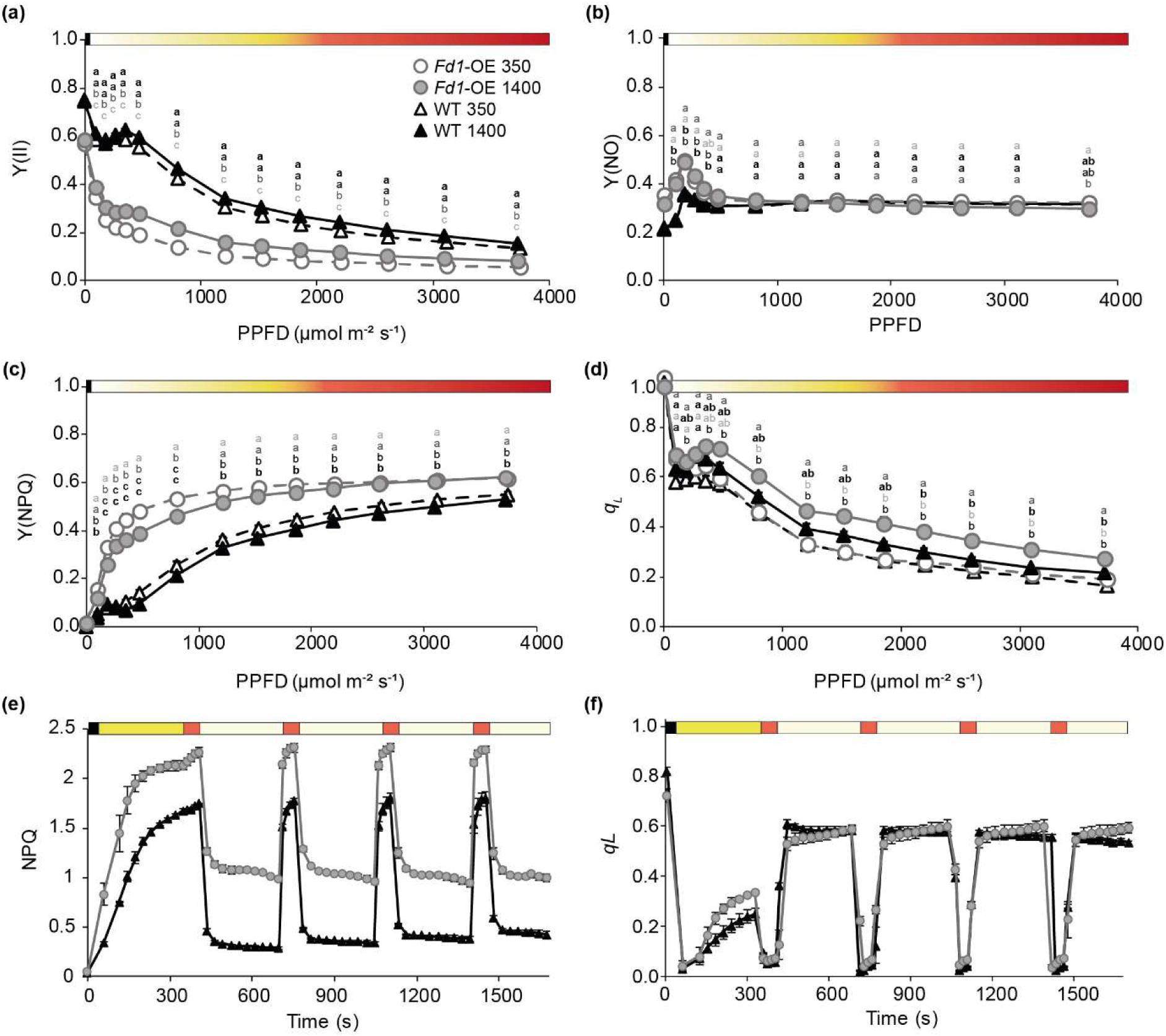
*Fd1*-OE plants exhibited a differential regulation of photosynthesis during rapid changes in light intensity as assessed by rapid light curves (RLC) and “fluctuating light” (FL) measurements. The responses of the photosynthetic parameters Y(II) (a), Y(NO) (b), Y(NPQ) (c) and *qL* (d) to fast changes in PAR illumination measured through RLC.. The responses of the photosynthetic parameters NPQ (e), and *qL* (f) to FL treatment. For both measurements WT (black triangles) and *Fd1*-OE (grey circles) plants were grown at 350 ((empty symbols and dashed lines) and 1400 μmol m⁻² s⁻¹ (filled symbols and solid lines). In FL treatment, yellow, white, and orange boxes indicates PPFD of 1400, 62 and 1961 μmol m⁻² s⁻¹. Data represent means ± SE (n = 10). Different letters indicate significant differences between WT and *Fd1*-OE plants within each PPFD (two-way ANOVA followed by Tukey’s multiple comparison test, p ≤ 0.05).

**Figure 7.**
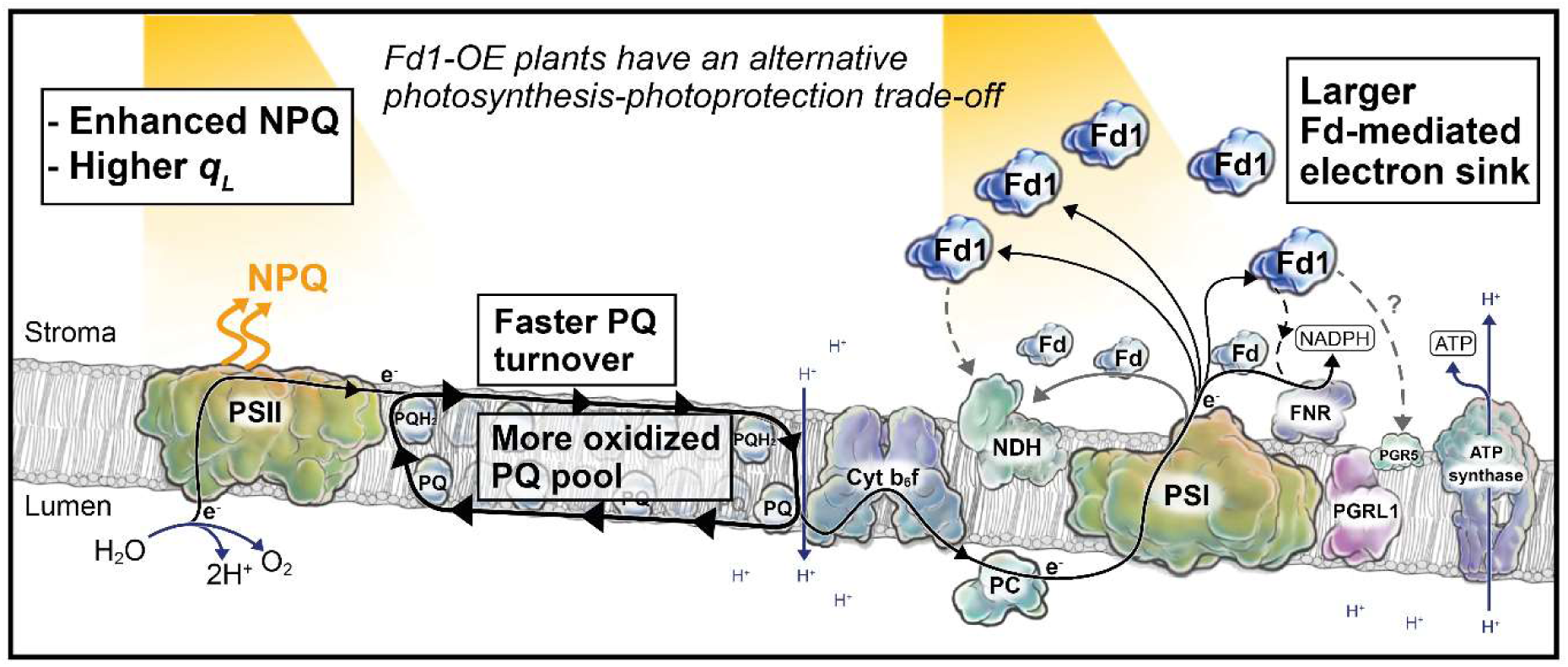
Proposed model of the alternative regulation of the photosynthesis in *Fd1*-OE plants. The coordinated adjustment between photochemical activity and energy dissipation establishes an alternative photosynthesis–photoprotection balance that improves adaptation to high irradiance and acclimation to fluctuating light conditions. Overexpression of heterologous Fd results in a faster plastoquinone (PQ) turnover underpinned by a larger electron sink downstream of PSI. This contributes to a more oxidized PQ pool and increased availability of open PSII reaction centers (higher *qL*), while maintaining elevated non-photochemical quenching (NPQ). Abbreviations: PSII Photosystem II, PQ Plastoquinone; PQH2 Plastoquinol; Cyt b6f; Cytochrome *b6f* complex; PC Plastocyanin; NDH NAD(P)H dehydrogenase-like complex; PSI Photosystem I; Fd Ferredoxin, Fd1 overxpressed (alien) ferredoxin; FNR Ferredoxin-NADP(+) oxidoreductase; PGR5 PROTON GRADIENT REGULATOR 5; PGRL1 PGR5-like protein-1.

The induction of photosynthesis in dark-adapted leaves was analyzed using fast chlorophyll a fluorescence induction (OJIP) curves (Fig. 5a) (Schansker *et al*., 2003; Schansker *et al*., 2005; Toth *et al*., 2007). Comparison of plants growing at 350 and 1400 μmol m⁻² s⁻¹ revealed marked differences across all phases of the induction curve. Under “high-light” growth conditions, differences were particularly evident in the O-J and J-I-P phases. During the O-J phase (photochemical phase, ∼2 ms), *Fd1*-OE plants displayed a steeper fluorescence rise than WT plants. This behavior has been associated with faster accumulation of reduced QA, the primary quinone electron acceptor of PSII (Schansker *et al*., 2003; Strasser *et al*., 2004; Toth *et al*., 2007). In contrast, *Fd1*-OE plants exhibited a further attenuated J-I phase, whose kinetics reflect the progressive reduction of the PQ pool between 20–30 ms after illumination (Schansker *et al*., 2005; Toth *et al*., 2007). Furthermore, *Fd1*-OE plants showed a delayed appearance of the P peak compared with WT (Fig. 5a, P and P*). To complement OJIP curve information that reflects changes in the coordination of the electron flow between PSII, the PQ pool and intersystem; PSI redox dynamics and the effective functional size of the PQ pool were further evaluated through absorbance changes around 830 nm (Fig. 5b) (Schreiber *et al*., 1988). P700 oxidation–reduction kinetics under far-red illumination revealed genotype-dependent differences that were maintained across growth irradiances, with *Fd1*-OE plants exhibiting a larger functional PQ pool size together with slower re-oxidation kinetics than their WT counterparts.

The fluorescence transient for transplastomic and control plants was also evaluated to study the response of photosynthesis to changes in illumination. The PIFR, or the rise in chlorophyll fluorescence upon switching off the actinic light on illuminated leaves during photosynthetic steady-state, has been used as a proxy of a reduction of the PQ pool by stromal reducing equivalents (Burrows *et al*., 1998; Gotoh *et al*., 2010). Both genotypes grown at 350 and 1400 μmol m⁻² s⁻¹ exhibited distinct PIFR traces during the light-to-dark transition. At both growth light intensities, *Fd1*-OE plants showed faster and stronger PIFR in comparison with control plants (Fig. 5c). Moreover, under “high-light” growth conditions, the post-illumination fluorescence rise exceeded the steady-state fluorescence level recorded immediately before actinic light was switched off.

In addition to dark-light transition analyses, RLCs were performed to assess the response of photosynthesis regulation in *Fd1*-OE plants to rapid increases in PAR intensity (Fig. 6a–d). Under both growth conditions, *Fd1*-OE plants displayed lower Y(II) and higher Y(NPQ) than WT plants across the dynamic illumination range, indicating a stronger engagement of regulated energy dissipation. No significant differences in passive energy dissipation (Y(NO)) were detected between genotypes across the PAR range between 500 and 3000 μmol m⁻² s⁻^1^, with only minor differences observed at the highest light intensities of the protocol (Fig. 6b). The largest differences in photochemical yield were observed around 350 μmol m⁻² s⁻^1^, where control plants reached maximal Y(II) (Fig. 6a). Above this irradiance, both Y(II) and Y(NPQ) progressively converged between genotypes. Interestingly, despite these differences in energy partitioning, and consistent with steady-state conditions, *qL* values remained significantly and consistently higher in *Fd1*-OE plants grown at 1400 μmol m⁻² s⁻^1^. Since *qL* reflects the fraction of open PSII reaction centers in relation to the redox poise of the PQ pool, these results indicate that *Fd1*-OE plants maintain a larger proportion of photochemically available centers across a broad dynamic range of irradiance. These results suggest *Fd1*-OE plants maintain higher photochemical availability with enhanced photoprotection during rapid changes in PAR when they grow at 1400 μmol m⁻² s⁻^1^.

The response of *Fd1*-OE plants grown at 1400 μmol m⁻² s⁻^1^ was further evaluated through a second set of PAM measurements using dynamic PAR intensities, also named fluctuating light treatments (FL) (Figs. 6e and f) (Tikkanen & Grebe, 2018; Yang *et al*., 2020). In our approach, plants were illuminated at their respective growth light intensity (yellow boxes), later challenged with brief periods of higher intensity PAR (photoprotection induction, orange boxes) followed by longer periods at lower PAR than growth irradiance (photoprotection relaxation, white boxes). Both *Fd1*-OE and control plants exhibited an NPQ induction in response to increases in PAR intensity which was sustained along the whole protocol (four high PAR challenges, orange boxes) (Fig. 6e). As observed in steady-state measurements, *Fd1*-OE plants grown at 1400 μmol m⁻² s⁻^1^ exhibited an augmented NPQ induction in absolute terms with similar kinetics during increases in PAR intensity. As previously observed in RLC, the *qL* parameter in *Fd1*-OE plants was also higher than in control plants during the initial growth light intensity period (Fig. 6f, yellow boxes). Moreover, after four cycles of PAR fluctuations, *Fd1*-OE plants sustained stable *qL* values, while WT plants showed a trend of decreasing values.

Together, OJIP, RLC, and FL analyses indicate that the improved performance of *Fd1*-OE plants under high-light growth conditions is accompanied by distinct changes in photosynthetic regulation. *Fd1*-OE plants grown at 1400 μmol m⁻² s⁻¹ consistently exhibited higher *qL* together with altered PQ pool-associated kinetics. These observations support the notion that elevated Fd1 levels modify the regulation of photosynthetic electron transport.

## Discussion

The balance between photochemical and photoprotective reactions determines the maintenance of photosynthetic performance under non-optimal growth conditions. Disruption of this balance often results in impaired photosynthetic electron transport, growth penalties, and permanent damage to sensitive components of the PETC, particularly PSI under sustained acceptor-side limitation (Lima-Melo *et al*., 2021; Tiwari *et al*., 2024). Accordingly, mutants affected in the coordination of plastoquinone (PQ) turnover and PSI electron flow (e.g. *immutans*, *pds3*, and PGR5-overexpressing plants) often exhibit photobleaching or leaf variegation as visible manifestations of this imbalance (Okegawa *et al*., 2007; Rosso *et al*., 2009; Okegawa *et al*., 2010). Previously, we characterized *Fd1*-OE plants, which also displayed a variegated phenotype associated with enhanced NPQ and growth penalties under greenhouse conditions (Blanco et al., 2013), suggesting an altered photosynthesis–photoprotection trade-off. Here, we demonstrate that increasing growth light progressively restores growth, photosynthetic performance, and leaf phenotype in *Fd1*-OE plants. This unexpected response indicates that elevated Fd1 levels redefine the boundary between permissive and non-permissive growth conditions, contrasting with the behavior of classical variegated mutants. Analysis of photosynthetic performance under steady-state and dynamic light regimes, together with chloroplast ultrastructure, revealed that this acclimation is associated with accelerated PQ turnover, a more oxidized PQ pool, and enhanced intersystem electron transfer. These changes are consistent with an increased Fd1-mediated electron sink and suggest that elevated Fd1 improves the coordination between photochemistry and photoprotection, enabling enhanced photoprotective capacity while maintaining efficient photosynthetic electron transport.

### A new photochemical-photoprotection trade-off in Fd1-*OE* plants improves higher growth light intensity adaptation

The first evidence for an altered photosynthesis-photoprotection trade-off in *Fd1*-OE plants comes from the failure to ameliorate growth penalties and reverse variegation by either reducing growth light intensity or supplying exogenous sucrose (Fig. 1). This observation indicates that the phenotype cannot be explained solely by carbon availability and instead points to an altered regulation of photosynthetic energy partitioning. Although Y(II) increased under lower irradiance, *Fd1*-OE plants continued to dissipate nearly 50% of absorbed light energy through non-photochemical processes (Klughammer & Schreiber, 2008). This reduced Y(II) under “greenhouse” (150-350 μmol m⁻² s⁻^1^) and “low-light” conditions (30 μmol m⁻² s⁻^1^) directly affected CO2 assimilation rate in *Fd1*-OE plants (Fig. 1c and Fig. S1c). Accordingly, *Fd1*-OE plants exhibited a lower initial slope of the ACO₂ light-response curve (A/PAR curve), reflecting a reduced apparent quantum yield (AQY) (Ogren, 1993). Together, these observations indicate that enhanced non-photochemical energy dissipation constrains photochemical efficiency and carbon assimilation under “low-light and “greenhouse” conditions, establishing these conditions as non-permissive for *Fd1*-OE growth. Therefore, *Fd1*-OE plants exhibit a photosynthetic regulatory behavior distinct from that of classical variegated mutants, in which increasing irradiance typically exacerbates rather than alleviates the phenotype (Rosso et al., 2009; Okegawa et al., 2010).

The disappearance of the variegated phenotype and the recovery of near-WT growth rates at 1400 μmol m⁻² s⁻¹ confirm the altered photosynthesis–photoprotection trade-off and a shift in the permissive growth limits towards higher light intensities in *Fd1*-OE plants (Fig. 2, Fig. S2). *Fd1*-OE plants showed an increase in pigments, improved CO2 assimilation rates and maximum quantum yield *Fv*/*Fm*, as well as a differential absorbed energy partitioning compared with control lines (Figs. 2 and 4, Tables 2 and 3). The phenotypical changes along the different growth light conditions were followed by a vast rearrangement of the thylakoid membranes in *Fd1*-OE plants, suggesting not a reversion to the WT phenotype but a more complex adjustment of the adaptive response (Fig. 3). The thylakoid rearrangements, characterized by strongly reduced grana stacking and increased numbers of enlarged plastoglobuli at the highest growth light intensities, partially resemble sun-type chloroplast acclimation responses (Fig. 3) (Lichtenthaler et al., 1981; Knapp & Smith, 1987). However, neither increased endogenous Fd content nor enrichment in minor Fd isoforms has been reported in these acclimation responses. Likewise, we did not detect changes in the abundance of endogenous or heterologous Fd, nor in other photosynthetic core proteins in *Fd1*-OE plants (Fig. 4a). Therefore, we hypothesize that this complex change in the adaptive response of *Fd1*-OE plants is underpinned by an augmented Fd1-based modification of the photosynthesis regulation.

Across the entire set of growth conditions, *Fd1*-OE plants simultaneously sustained higher NPQ and *qL* relative to control plants (Table 3, Figs. 4c, S3 and S4). Higher *qL* values indicate a higher fraction of open reaction centers at PSII capable of performing photochemistry in *Fd1*-OE plants (Baker, 2008). Specifically, the *qL* parameter is influenced both by excitation energy transfer to PSII reaction centers and by the redox poise of the PQ pool, which is supported by the presence of the F₀′ parameter in its formula. Therefore, the simultaneously higher values in NPQ and *qL* suggest a coordinated adjustment between dissipative and photochemical processes in *Fd1*-OE plants. In this context, the sustained NPQ together with the larger fraction of open PSII centers is consistent with a more oxidized PQ pool and a coordinated regulation of photochemical and dissipative processes under increasing irradiance. Indeed, *Fd1*-OE plants exhibited a more balanced variation in the fractions of absorbed energy used for photochemistry (Y(II)) and dissipated by heat (Y(NPQ)) between non-permissive and permissive growth conditions (350 and 1400 µmol m⁻² s⁻¹) relative to control plants (Table 3). Aligned with this interpretation, the lower Y(NO) values observed in *Fd1*-OE plants grown at “high-light” intensities indicate a more optimized energy allocation (Table 3), and a concomitant improvement in AQY derived from A/PAR curves supporting the increased growth rates (Figs. 2 and 4b; Table 3). Altogether, these results indicate that the adaptive response of *Fd1*-OE plants to high growth irradiance is closely associated with the redox poise and turnover of the PQ pool. The simultaneous maintenance of high NPQ and a larger fraction of open PSII reaction centers suggests a coordinated regulation between dissipative and photochemical processes, likely sustained by an enhanced electron sink downstream of the PQ pool.

### Mechanisms underlying the new photochemical-photoprotection trade-off in Fd1-OE plants and its impact on acclimation response

While PSII steady-state photosynthetic parameters support a central role for PQ turnover in the coordinated adjustment to “high-light” conditions, no major differences were detected in parameters associated with PSI functioning, where heterologous Fd1 is expected to operate in *Fd1*-OE plants (Table 3). Previous studies of lines overexpressing other PSI-associated proteins, such as PGR5 and Fd, also reported substantial alterations in photosynthetic regulation but comparatively minor phenotypic consequences and differences in PSI steady-state parameters (Yamamoto et al., 2006; Okegawa et al., 2007). The general phenotypic changes, and specifically thylakoid rearrangements, including LHC and pigments organization, may be in part offsetting effect on steady-state PSI parameters in *Fd1*-OE plants (Table 1–3).

These limitations were overcome by challenging photosynthetic performance under changing light conditions, where the kinetics of PQ reduction and re-oxidation can be more directly resolved (Figs. 5 and 6). To address the mechanisms underlying the altered PQ redox poise during photosynthesis induction, we analyzed the polyphasic chlorophyll fluorescence transient rise (OJIP curves) in dark-adapted plants (Fig. 5a). Because the successive OJIP phases reflect sequential restrictions in electron flow along the intersystem of the PETC (Schansker et al., 2005), the fluorescence kinetics of *Fd1*-OE plants provide additional evidence for a faster redox turnover of the PQ pool. The J phase mainly reflects the reduction and accumulation of the primary PSII acceptor QA, which was more pronounced in *Fd1*-OE plants than in control plants (Toth et al., 2007). In contrast, the subsequent J-I phase, associated with PQ turnover kinetics, was strongly dampened in *Fd1*-OE plants, and even developed a transient depression in plants grown at 1400 µmol m⁻² s⁻¹, instead of the sustained J-I rise observed in control plants (Schansker et al., 2005; Toth et al., 2007). Altogether, these kinetics indicate faster redox turnover rates at both the donor and acceptor sides of the PQ pool in *Fd1*-OE plants.

Furthermore, the delayed appearance of the P peak (P*) in *Fd1*-OE plants is consistent with an enhanced electron sink at the PSI acceptor side, supporting the accelerated PQ turnover. Accordingly, the overall kinetics of photosynthesis induction suggest that *Fd1*-OE plants sustain faster PQ pool turnover through an increased withdrawal of electrons downstream of PSI. Complementary evidence supporting this conclusion was obtained from kinetics measurements of PSI redox on P700 absorbance (ΔA₈₂₀) (Fig 5c). In particular, *Fd1*-OE plants exhibited a relatively larger effective functional PQ pool size, which is aligned with the idea of an enhanced PQ turnover driven by an enlarged electron sink at the PSI acceptor side, and the PQ re-reduction kinetics (see below) (Schansker et al., 2003; Klughammer, C. & Schreiber, U, 2008).

The participation of Fd1 in the downstream electron sink underlying photosynthetic regulation in *Fd1*-OE plants was further supported using a second dynamic-light approach based on post-illumination fluorescence rise (PIFR) kinetics (Fig. 5b). This transient increase in fluorescence, originally described as the F0 rise, has been extensively associated with the re-reduction of the PQ pool and its equilibration with QA and QB at PSII through stromal reducing equivalents after illumination is switched off (Burrows *et al*., 1998; Gotoh *et al*., 2010). Compared with control plants, *Fd1*-OE plants displayed a stronger and more prolonged fluorescence rise, reaching higher fluorescence levels in darkness, particularly under 1400 µmol m⁻² s⁻¹ growth conditions (Fig. 5b, right panel). Considering the higher *qL* values and faster PQ turnover in *Fd1*-OE plants, PIFR kinetics suggest a larger or more rapidly mobilized pool of stromal reducing equivalents capable of re-reducing the PQ pool. Moreover, the sustained PIFR signal in *Fd1*-OE plants differs than of other mutant signals characterized by an augmented pool of stromal reducing equivalents, such as *fba3-1* and *pgdh3-1* (Gotoh et al., 2010; Kramer et al., 2024). suggesting a more direct participation of Fd1 in PQ re-reduction. Although PIFR signals have been associated with PSI-cyclic electron transport pathways involving NDH and, more recently, PGR5-dependent processes (Burrows et al., 1998; Sazanov et al., 1998; Gotoh et al., 2010; Kramer et al., 2024), the present results do not allow assignment to a specific pathway or confirm a mechanistic involvement. Nevertheless, the enhanced PIFR response in *Fd1*-OE plants provides additional evidence supporting an active participation of Fd1 in regulating electron flow downstream of PSI. This dynamic cycling of reducing equivalents back into the PQ pool may also contribute to the slower PSI re-oxidation kinetics observed in *Fd1*-OE plants during dark transitions (Fig 5c).

The emerging picture of Fd1 acting as an enhanced downstream electron sink, sustaining high rates of PQ turnover together with elevated NPQ levels confirms the existence of a differential regulation of photosynthesis in *Fd1*-OE plants. The final set of experiments provides a proof-of-concept for this model by challenging plants under highly dynamic illumination regimes. Measurements of *qL* and NPQ during rapid stepwise increases in irradiance (RLC) and fluctuating light conditions showed that *Fd1*-OE plants sustain more stable *qL* values across contrasting illumination patterns (Figs. 6d and f).

The integration of these results with the broader physiological characterization indicates that Fd1 overexpression establishes an alternative balance between photochemical and photoprotective processes, underpinning both improved adaptation to high growth irradiance and enhanced acclimation capacity under changing environmental conditions. Altogether, these findings demonstrate that engineering electron partitioning downstream of PSI by Fd may be part of novel strategies to optimize photosynthetic performance and resilience under naturally fluctuating environments.

## Acknowledgments

This work was supported by grants from ANPCyT (PICT-2020-SERIEA-01326, PICT-2021-I-A-00373 and PICT-2021-CAT-II-00042) and the MSCA-EPPN Project “E.A.S.Y.”. N.E.B. is a researcher of the Argentine National Scientific and Technical Research Council (CONICET). J.G.B. and C.B. were supported by fellowships associated with the PICT projects, and C.B. is currently a CONICET fellow.

We kindly thank Mingwei Geng for assistance with pigment determinations. We also thank Carl-Otto Ottosen (Aarhus University, Denmark, EPPN Project host) and the staff of the Dynapheno Platform for their excellent technical support. Finally, we thank Eva Rosenqvist for critical comments.

## Author contributions

NEB conceived the project and experiments, as well as performed first set of experiments. CL and NEB made low light experiments. JGB performed most of the PAM and P700 measurements (steady-state and under changes in light intensity), pigment determination, Western blots, and physiological determination. MM was responsible of ultrastructure TEM microscopy. NEB and JGB made the main analysis, draft conception, and first round of draft writing. All authors contribute to data analysis, draft writing and critical reading. All authors read and approved the manuscript.

## Data availability

Data is included in Supplementary information. Any data is also available upon request to.

## Supporting information

**Figure S1.**
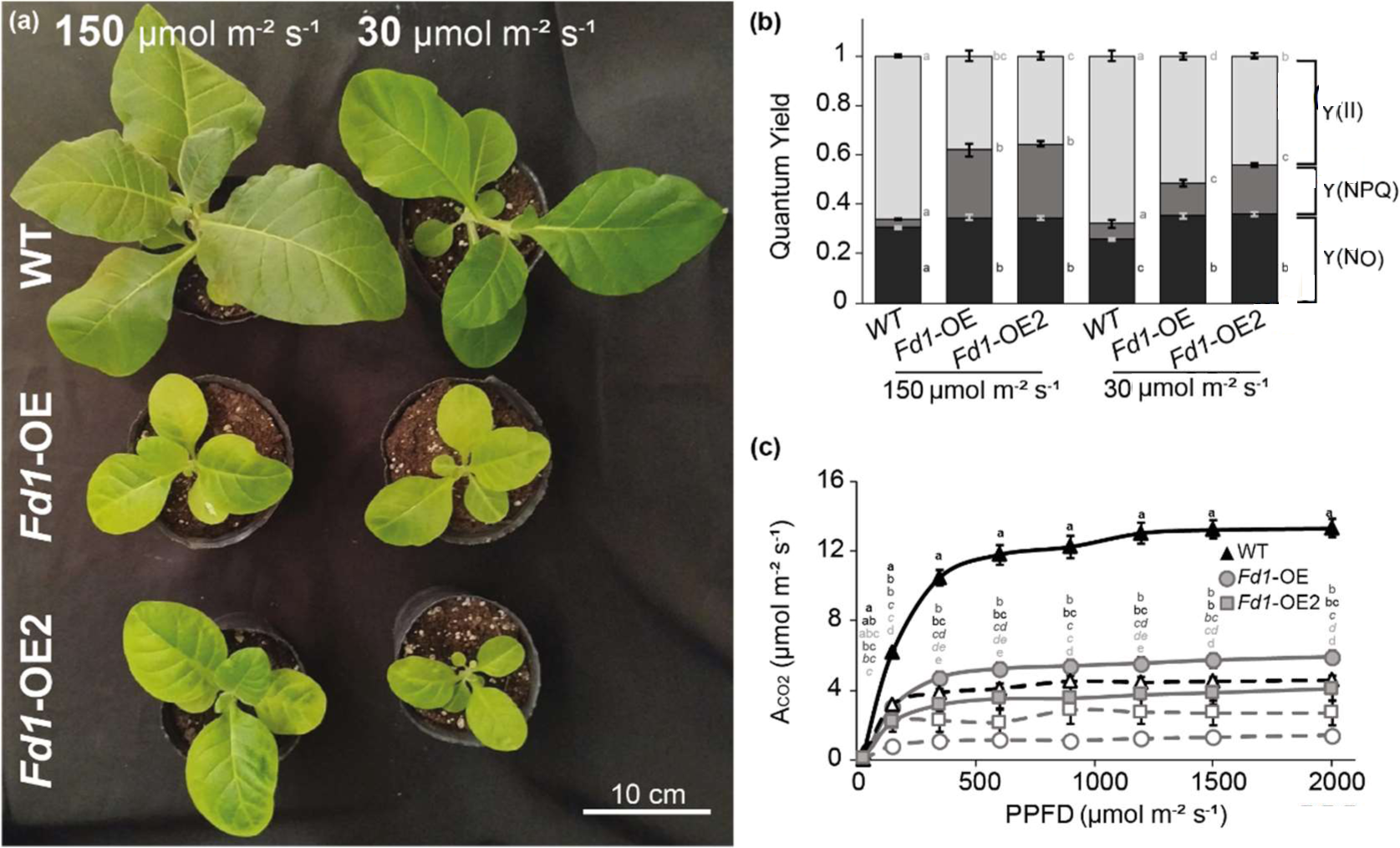
Phenotypic evaluation of independent transplastomic lines expressing *Fd1* construct. (a) 5-week-old control (WT, top) and two ferredoxin overexpressing lines (*Fd1*-OE, middle and *Fd1*-OE2, bottom) plants grown under “greenhouse” (150 µmol m-2 s-1) and “low light” (30 µmol m-2 s-1) conditions. (b) Steady-state PS II quantum yield parameters Y(II) (light grey), Y(NPQ) (dark grey) and Y(NO) (black), measured at indicated growth light. (c) Light-response curves of CO₂ assimilation rates for WT (black triangles), *Fd1*-OE (grey circles) and *Fd1*-OE2 (grey squares) plants. Dashed and empty symbols, and solid lines and filled symbols correspond to “low-light” and “greenhouse” conditions, respectively. Data points represent the means ± SE (n-10). Different letters indicate statistical significance between genotypes at each light intensity, (two-way ANOVA followed by Tukeýs multiple comparison test, p ≤ 0.05).

**Figure S2.**
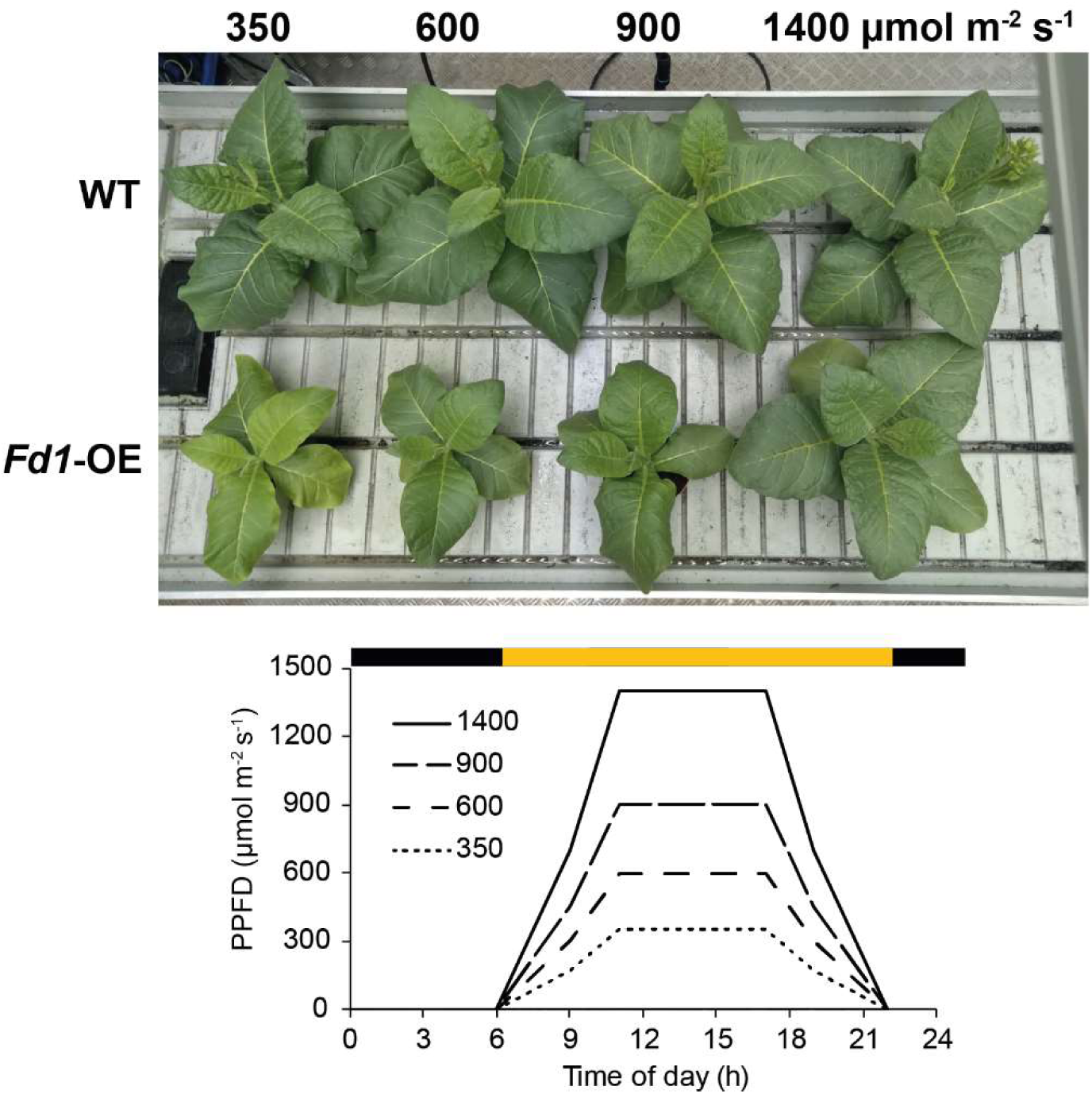
Outdoor-like dynamic light regimes used to evaluate light acclimation responses across contrasting growth irradiances. Plants were cultivated under a 16/8 h light–dark photoperiod with gradual ramping at both light onset and offset to mimic outdoor conditions. Light intensity increased in two consecutive phases, reaching 50% of the maximum intensity within the first 3 h and the full peak after an additional 2 h, followed by 6 h at maximum irradiance. An analogous ramping profile was applied during the transition to darkness. Representative images of WT and *Fd1*-OE plants grown at 350, 600, 900 and 1400 µmol m-² s-¹ are shown.

**Figure S3.**
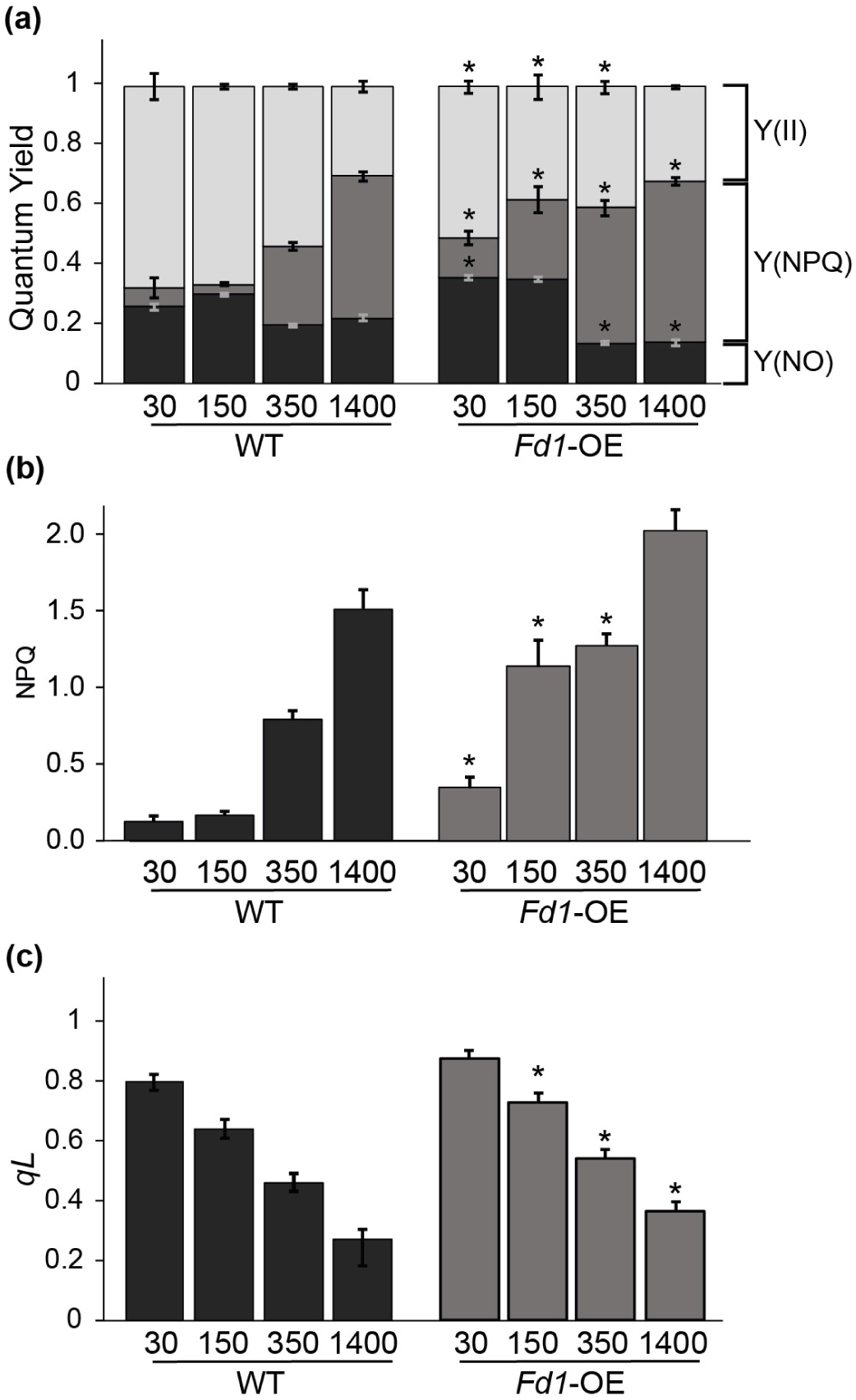
Steady-state parameters for plants grown across a wide range of light conditions. Chlorophyll fluorescence measurements of dark-adapted and steady-state parameters were performed after 20 minutes of darkness and 12 minutes of growth light intensity exposure, respectively. Measurements were performed on the first fully expanded leaf of 5-week-old plants. Data represent means ± SE (n ≥ 10) for each condition. Asterisks (*) indicate significant differences relative to WT plants grown under the same conditions (one-way ANOVA followed by Tukey’s multiple comparison, p ≤ 0.05).

**Table S1.**
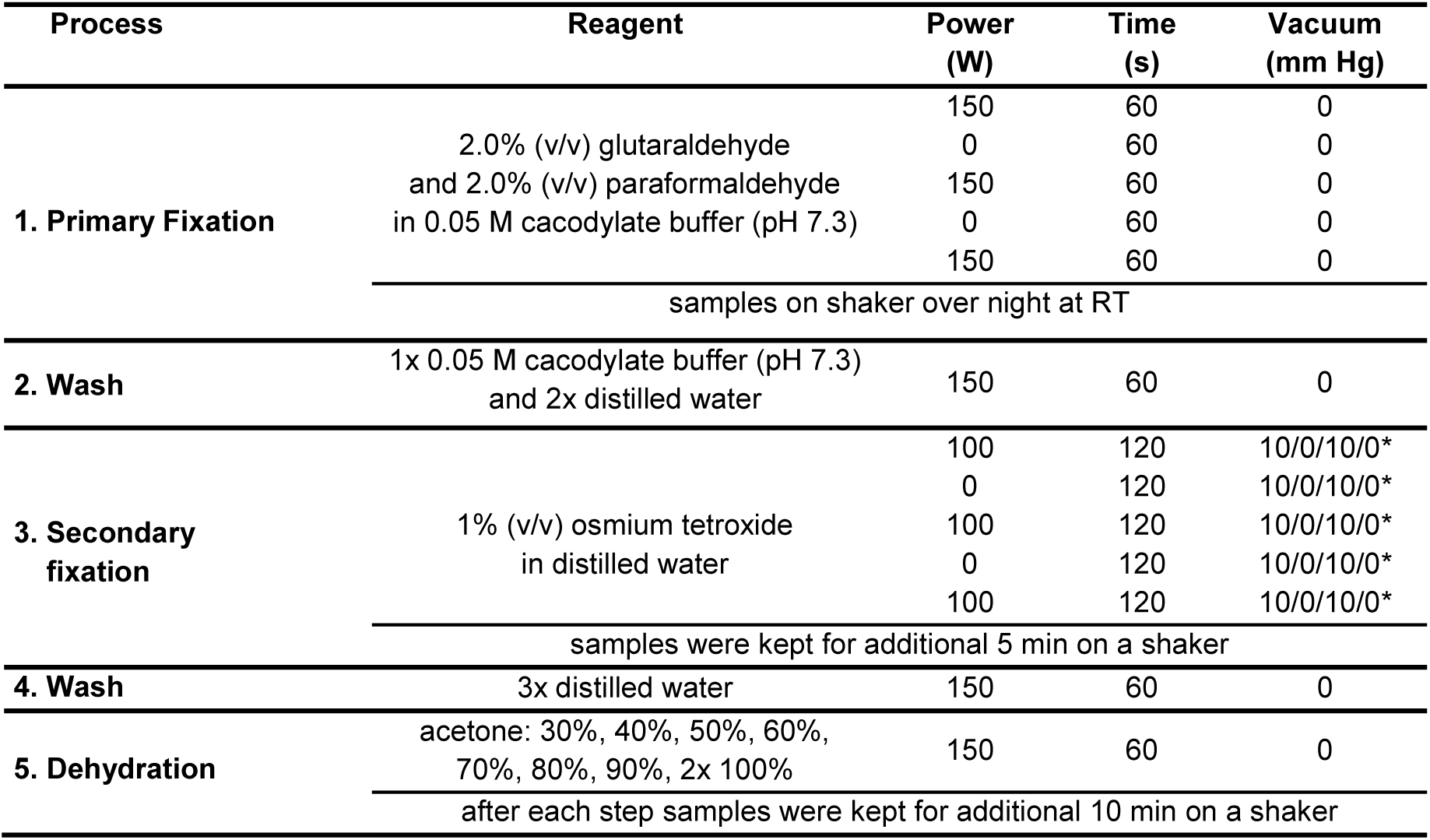
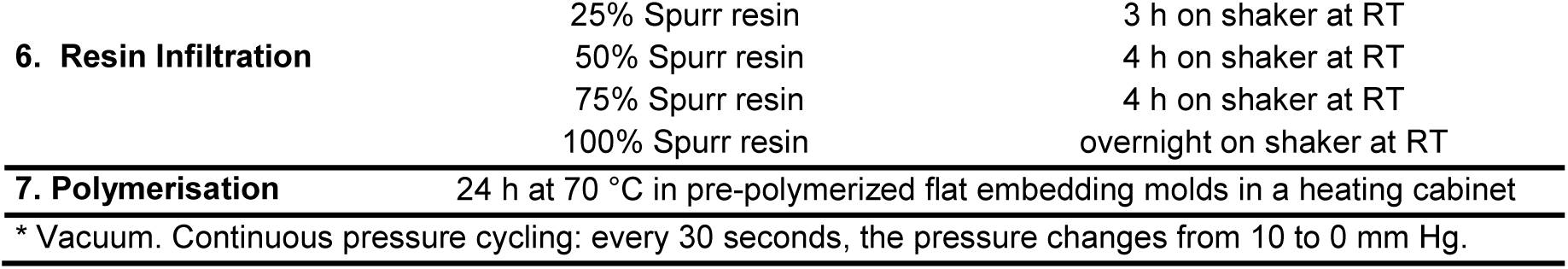
Combined conventional & microwave assisted plant tissue preparation in a PELCO Bio Wave 34700-230 (Ted Pella, Inc., Redding CA, USA)

| Process | Reagent | Power (W) | Time (s) | Vacuum (mm Hg) |
| --- | --- | --- | --- | --- |
| 1. Primary Fixation | 2.0% (v/v) glutaraldehyde and 2.0% (v/v) paraformaldehyde in 0.05 M cacodylate buffer (pH 7.3) | 150 | 60 | 0 |
|  |  | 0 | 60 | 0 |
|  |  | 150 | 60 | 0 |
|  |  | 0 | 60 | 0 |
|  |  | 150 | 60 | 0 |
|  | samples on shaker over night at RT |  |  |  |
| 2. Wash | 1x 0.05 M cacodylate buffer (pH 7.3) and 2x distilled water | 150 | 60 | 0 |
| 3. Secondary fixation | 1% (v/v) osmium tetroxide in distilled water | 100 | 120 | 10/0/10/0* |
|  |  | 0 | 120 | 10/0/10/0* |
|  |  | 100 | 120 | 10/0/10/0* |
|  |  | 0 | 120 | 10/0/10/0* |
|  |  | 100 | 120 | 10/0/10/0* |
|  | samples were kept for additional 5 min on a shaker |  |  |  |
| 4. Wash | 3x distilled water | 150 | 60 | 0 |
| 5. Dehydration | acetone: 30%, 40%, 50%, 60%, 70%, 80%, 90%, 2x 100% | 150 | 60 | 0 |
|  | after each step samples were kept for additional 10 min on a shaker |  |  |  |

|  |  |  |
| --- | --- | --- |
| 6. Resin Infiltration | 25% Spurr resin | 3 h on shaker at RT |
|  | 50% Spurr resin | 4 h on shaker at RT |
|  | 75% Spurr resin | 4 h on shaker at RT |
|  | 100% Spurr resin | overnight on shaker at RT |
| 7. Polymerisation | 24 h at 70 °C in pre-polymerized flat embedding molds in a heating cabinet |  |
| * Vacuum. Continuous pressure cycling: every 30 seconds, the pressure changes from 10 to 0 mm Hg. |  |  |

